# Investigation of normalization procedures for transcriptome profiles of compounds oriented toward practical study design

**DOI:** 10.1101/2023.10.01.560398

**Authors:** Tadahaya Mizuno, Hiroyuki Kusuhara

## Abstract

The transcriptome profile is a representative phenotype-based descriptor of compounds, widely acknowledged for its ability to effectively capture compound effects. However, the presence of batch differences is inevitable. Despite the existence of sophisticated statistical methods, many of them presume a substantial sample size. How should we design a transcriptome analysis to obtain robust compound profiles, particularly in the context of small datasets frequently encountered in practical scenarios? This study addresses this question by investigating the normalization procedures for transcriptome profiles, focusing on the baseline distribution employed in deriving biological responses as profiles. Firstly, we investigated two large GeneChip datasets, comparing the impact of different normalization procedures. Through an evaluation of the similarity between response profiles of biological replicates within each dataset and the similarity between response profiles of the same compound across datasets, we revealed that the baseline distribution defined by all samples within each batch under batch-corrected condition is a good choice for large datasets. Subsequently, we conducted a simulation to explore the influence of the number of control samples on the robustness of response profiles across datasets. The results offer insights into determining the suitable quantity of control samples for diminutive datasets. It is crucial to acknowledge that these conclusions stem from constrained datasets. Nevertheless, we believe that this study enhances our understanding of how to effectively leverage transcriptome profiles of compounds and promotes the accumulation of essential knowledge for the practical application of such profiles.

## Introduction

In order to utilize in silico analysis for understanding the properties of a compound such as toxicity, it is first necessary to describe the compound as numeric information. The most frequently used methodology is structure-based description, and various descriptors have been developed (Rogers and Hahn, 2010; Duvenaud *et al*., 2015; Le *et al*., 2020; Jaeger *et al*., 2018; Sawada *et al*., 2014; Nemoto *et al*., 2023). In recent years, the phenotype-based descriptors have been utilized as well, which are based on the phenotypes of cultured cells or animals subjected to the compound (Sutherland *et al*., 2019; Igarashi *et al*., 2015; Chandrasekaran *et al*., 2021; Waters *et al*., 2008). Phenotype-based descriptors are superior in that they are structure-independent, allowing evaluation of mixtures and other cases in which the structure is not uniquely determined, and in that they account for association with biological systems to several degree.

On the other hand, unavoidable issues exist regarding the phenotype-based descriptors since data must be obtained in an experimental manner. Transcriptome-based profiling is one of the major methodologies in phenotype-based descriptors and large databases exist such as connectivity map (CMap) and L1000, which are widely accepted as data sources for in silico analysis (Lamb, Emily D. Crawford, et al., 2006; Subramanian et al., 2017). In addition to unavoidable cost for data acquisition, a major problem of these approaches is batch differences such as inter-trial difference and center difference (Nygaard *et al*., 2016; Sutherland *et al*., 2016). Note a batch denotes a dataset comprising transcriptome data derived from a singular platform in this context. Lots of researchers have tried to address the problem from various angles such as usage of spike-in control in the data acquisition step and statistical correction in the data processing step (Luo *et al*., 2010; Sun *et al*., 2011; Müller *et al*., 2016; Haghverdi *et al*., 2018). Established and widely recognized techniques, such as Combat, address batch disfferences among transcriptome profiles through empirical Bayes methodology, as outlined by Johnson et al. and Leek et al. (Johnson *et al*., 2007; Leek *et al*., 2012). However, it is imperative to exercise caution in the statistical application of Combat, particularly concerning the parametric nature of the prior. Consequently, how to overcome batch differences in transcriptome profiles remains to be an open question in the field including profiles for compounds.

Gene expression data obtained with transcriptome analysis are relatively conserved in the baseline expression levels of the target (e.g., *Alb* gene coding albumin is highly expressed in liver). Thus, such *expression profiles* are not suitable as descriptors of compound characteristics without appropriate gene-wise normalization. Particularly, transcriptome profiles of compounds should be normalized against control treatment such as DMSO and PBS treatments for utilization of existing knowledge of compounds, which we call *response profiles* in this study (Lamb, Emily D Crawford, et al., 2006; Subramanian et al., 2017; Mizuno et al., 2019, 2020). With respect to its practical utilization as a query for large transcriptome profile databases like CMap and LINCS, few knowledge exists regarding normalization methodologies for ensuring the robustness of compound transcriptome profiles against batch differences, particularly in the context of small datasets. In this study, we examined normalization procedures for transcriptome profiles of compounds focusing on methods (how to select the baseline and to conduct normalization) and, in the first place, study design for acquiring GeneChip (microarray) data (how many control samples should be prepared). First, in order to obtain knowledge on the normalization procedures, we examined it with a large dataset with a sufficient data size. Next, setting the optimized large dataset as ground truth, we evaluated the study design for a small dataset. We addressed these points based on the similarity of transcriptome profiles of the same compound among different data sets.

Regarding the large dataset of transcriptome profiles of compounds, the study by Iskar et al. using the CMap dataset, as well as the present study, provides important insights in this field and several groups follow them (Iskar *et al*., 2010; Iwata *et al*., 2022; Fernández-Torras *et al*., 2022). However, they examined methodologies that do not take deviation into account and the impact of the batch correction methods is not examined. The iLINCS portal, which compiles data from the CMap and LINCS projects, provides the processed transcriptome profiles and carefully discloses the processing procedures they use (Subramanian et al., 2017; Lamb, Emily D Crawford, et al., 2006; Pilarczyk et al., 2022). While it is informative, the process of selection is not clear, and we cannot gain insight into which process was important. Finally, considering stability of deviation, normalization procedures are different between the large dataset such as CMap and small dataset in practical use, but there is no study on the processing procedures for transcriptome profiles of compounds handling small-scale data, to the best of our knowledge. It should be noted that this study deals with microarray data, and both results and discussions are grounded in microarray outcomes, despite RNA-seq being the primary platform for transcriptome profiling. However, it is also noteworthy that in terms of the multivariate properties delineating compound effects, microarray data and RNA-seq data exhibit considerable similarity (Wang *et al*., 2014; Xu *et al*., 2013; Zhao *et al*., 2014). Taking into account the legacy data accumulation, scale considerations, batch differences, and the specific objectives of this study, microarray data sourced from CMap was employed.

## Materials and Methods

### 1. Data preparation

#### 1.1. Dataset

In this study, we employed microarray data obtained by the CMap project build 2 (https://clue.io/data/CMB02#B02). Microarray data and meta information are available from the web page of the project and then is subject to processing described below. The content is shown in **Table 1**. In this study, we manage two datasets, namely HT-HG-U133A and HT-HG-U133A_EA, representing distinct arrays from Affymetrix, with evident differences in batches. Note that each dataset is composed of data derived from several batches and the number of samples in each batch is shown as histograms in **Supplementary Figure 1a**. It is also noteworthy that these arrays consist of nearly identical gene sets, exhibiting only a marginal percentage difference and similarity.

**Table 1.**
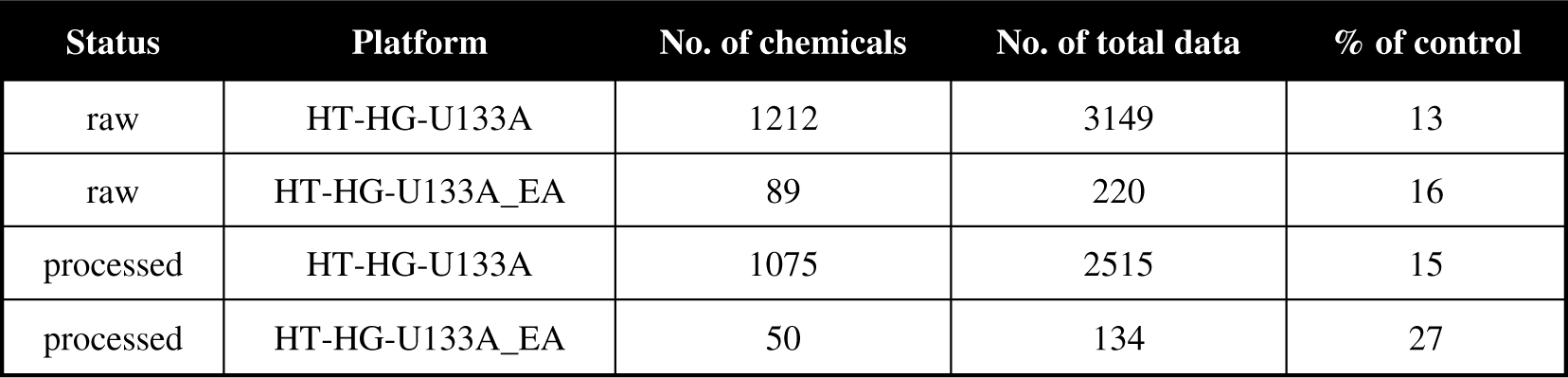
Information of data employed in this study.

| Status | Platform | No. of chemicals | No. of total data | % of control |
| --- | --- | --- | --- | --- |
| raw | HT-HG-U133A | 1212 | 3149 | 13 |
| raw | HT-HG-U133A_EA | 89 | 220 | 16 |
| processed | HT-HG-U133A | 1075 | 2515 | 15 |
| processed | HT-HG-U133A_EA | 50 | 134 | 27 |

#### 1.2. Preprocessing

Microarray data is provided as CEL files of Affymetrix platform. The downloaded files were subjected to *affy* package in Bioconductor library of R (version 3.1.6) and converted into gene expression data with MAS5 in default setting. Expression data was rearranged by cell lines and data of MCF7 cells was processed in further steps. Note that data derived from MCF7 cells was chosen to ensure an adequate sample size and batch quantity.

### 2. Processing procedures

As in prior works such as CMap, transcriptome profiles of compounds are defined as the profiles of gene expression differences from the indicated control and we call it response profiles in this study. Here, we describe the details of converting expression profiles to response profiles. The flow of processing procedures is summarized in **Supplementary Figure 1b** and we wrapped them as *exp2res* package in python 3.

#### 2.1. Imputation and trimming

Expression data was rearranged by cell lines and data of MCF7 cells was processed in further steps. NaN values of raw expression data were imputed with the mean value in gene-wise if 80% of samples exhibited not NaN and otherwise the genes including NaN were removed. Genes scored less than 1.0 were also removed.

#### 2.2. Log2 conversion, ID conversion, and index summarization

Trimmed data was subjected to log2 conversion. Then, probe IDs were converted to gene symbol and the median of value with the same symbol was taken as the representative expression of the corresponding gene.

#### 2.3. Batch correction and quantile normalization

Based on batch information given in the meta file, Combat was applied to correct batch information using *pyCombat* package (version 0.3.2, https://github.com/epigenelabs/pyComBat) in python with default setting and then quantile normalization was done.

#### 2.4. Difference calculation

We obtained response profiles of compounds as follows. Let *X ∈ ℝ^p×n^* be microarray data (expression data) of the cells treated with compounds, where p and n are the dimension of genes and the number of samples. Difference calculation operator *f* maps expression data *X* to response profiles of compounds *Y ∈ ℝ^p×n^*:

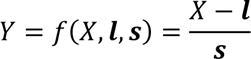

where *l ∈ ℝ^p^* and *s ∈ ℝ^p^* are location and scale parameters derived from control data and define the calculation methods: z, modified z, and robust z method. Note that we have two options as control data: only expression data of the cells treated with solvent (e.g., DMSO and PBS) and all expression data including both compounds of interest and solvent. In addition, considering batch difference, we also need to choose whether the above calculation is conducted within each batch or not (Iskar *et al*., 2010). We discuss details in **Results and Discussion**.

In z method, *l* and *s* is the mean and standard deviation values of control data in feature-wise, respectively. Thus, the method is z-normalization in feature-wise when all expression data is employed as control data while the intuitive interpretation of the outcome scores in the other case is an indicator of how much the data of the treatment group are outliers in the distribution of the controls assuming normal distribution. The concept of modified z and robust z method is the same and how to calculate *l* and *s* is different from those of z method. With respect to *l*, both methods employ the median values of control data in feature-wise while s values in modified z and robust z method are calculated as median absolute difference (MAD) and interquartile range (IQR) scaled to normal distribution:

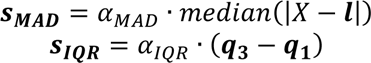

where *a_MAD_ = 1.4826* and *a_JQR_ = 0.7413* are the scaling parameter to normal distribution, respectively, and *q_3_* and *q_l_* are the 3^rd^ and 1^st^ quantile of control data in feature-wise, respectively.

#### 2.5. Quality check

In order to filter out raw quality data, we set the quality check step based on response profile consistency within biological replicates, which we call intra-dataset consistency. We assume that response profiles of biological replicates of a compound should be similar each other compared with those with the other compounds in the dataset.

Let *S = {d_k_|k = 1, ⋯, N}* be the dataset of interest where *N* is the total amount of data and *d_k_* corresponds to the response profile *y_k_*. Let us consider *S_i_*, a subset of *S* that is composed of biological replicates of a compound *i*. For each member of *S_i_*, we calculate correlation with the other response profiles considering whether the counterpart belongs to *S_i_* or not, and obtain variable *c_rep,k_* and *c_null,k_* that are the correlation of *y_k_* with the data in and not in *S_i_*, respectively. Based on the indicated threshold *r*, we check the quality of the focused response profile *y_k_*, corresponding to *d_k_*, by the following criterion:

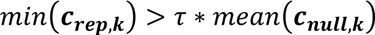

where *min(·)* is the minimum operator and *mean(·)* is the average operator. In this study, we investigated *r = {0.0, 0.5, 1.0, 1.5, 2.0, 2.5, 3.0}*.

#### 2.6. Consensus signature and sample summarization

Response profiles passing the quality check step are corrected with consensus signature. This method was first introduced to microarray data handling by LINCS project and employs information of biological replicates and corrects each response profile based on correlation within biological replicates as follows:

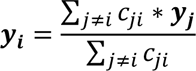

where *y_i_* is a response profile of the *i* th biological replicate of a compound and *c_ji_* is the correlation between the *i*th and *j*th profiles in biological replicates of a compound. The corrected profiles of are summarized with averaging in feature-wise within each compound to complete the processing flow.

### 3. Metrics

#### 3.1. Intra-dataset consistency

We define intra-dataset consistency as a metric to investigate quality of response profiles of compounds under the same assumption in the quality check step: robustness within a dataset. This metric is calculated as distance between the empirical distribution of correlation values within biological replicates *F_rep_(x)* and the empirical distribution of those that are not *F_null_(x)* and given as Kolmogorov–Smirnov (KS) statistic:

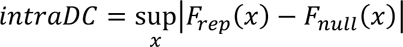

where 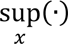 is the supremum operator. The statistic was calculated with *scipy.stats* module in python 3.

#### 3.2. Inter-dataset consistency

As with intra-dataset consistency, we assume that correlation of response profiles of the same compound is high as well even across datasets if the quality of profiles is high. Let *F_match_(x)* be the empirical distribution of correlation values of response profiles of the same compound pairs across datasets and *F_other_(x)* be the empirical distribution of those between other compounds. Inter-dataset consistency is also defined as KS statistic:

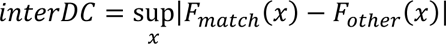

Note that inter-dataset consistency, robustness across datasets, is available if the two datasets to be analyzed have enough number of the same compounds. The size of the intersection of HT-HG-U133A and HT-HG-U133A_EA compounds is 22 and the member is listed in **Table 2**.

**Table 2.** List of compounds employed for investigating inter-dataset consistency.

| **Name** | **Description** |
| --- | --- |
| 15-delta prostaglandin J2 | prostaglandin J derivative |
| acetylsalicylic acid | non-steroidal antiinflammatory agent |
| alpha-estradiol | estrogen |
| alvespimycin | HSP90 inhibitor |
| estradiol | estrogen |
| felodipine | calcium channel blocker |
| fulvestrant | steroidal antiestrogen |
| geldanamycin | HSP90 inhibitor |
| LY-294002 | PI3K/BET Inhibitor |
| monorden | HSP90 inhibitor |
| nordihydroguaiaretic acid | potent lipoxygenase inhibitor |
| prochlorperazine | phenothiazine |
| pyrvinium | quinolinium ion |
| resveratrol | phytoalexin |
| sirolimus | mTOR inhibitor |
| tanespimycin | HSP90 inhibitor |
| thioridazine | phenothiazine |
| tretinoin | all-trans-retinoic acid |
| trichostatin A | HDAC inhibitor |
| trifluoperazine | phenothiazine |
| vorinostat | HDAC inhibitor |
| wortmannin | PI3K inhibitor |

### 4. Investigation of the relationship between the number of control samples for response profile calculation and inter-dataset consistency

Using the response profiles from the extensive dataset (HT-HG-U133A) as the ground truth for the response profiles of these compounds, we compared the response profiles of the smaller dataset, which were derived from randomly selected control samples. The smaller dataset (HT-HG-U133A_EA) encompasses a total of 36 control samples (DMSO). For the 22 compounds shared between both datasets, we applied batch correction using Combat to the smaller dataset. We then sampled the indicated number of control samples (greater than 2 for scale calculation) and obtained response profiles as MAD z with vehicle control. Note that the number of treatment samples are 2-3 and response profiles of a compound was aggregated with consensus signature in section 2.1. Subsequently, we assessed the inter-dataset consistency against the ground truth response profiles obtained from the larger dataset. Note that we employed MAD z with batch correction and considered all samples from the whole datasets for scale calculation. This procedure was iterated 50 times, and the resulting values were presented.

### 5. Data and code availability

Analysis was mainly performed using Python (version 3.9.12) with open-source packages (numpy, version 1.22.4; pandas, version 1.1.5; scipy, version 1.8.0; scikit-learn, version 1.0.2; statsmodels, version 0.13.2) unless otherwise noted. Code is available at https://github.com/mizuno-group/NormTranscriptome.

## Results and Discussion

### 1. Comparison of intra-dataset consistency of response profiles between conversion methods

Good response profiles for a compound of interest should exhibit a notable similarity among biological replicates within a dataset. Thus, we investigated the influence of calculation methodologies on the similarity of response profiles between biological replicates, referred to as intra-dataset consistency (intraDC) in this study. IntraDC is an indicator of robustness within a dataset. This analysis was conducted on two relatively large microarray dataset sourced from CMap (HT-HG-U133A and HT-HG-U133A_EA, as detailed in **Table 1**). The following aspects were compared: Firstly, the effect of employing batch correction versus not, considering that a majority of large datasets encompass a variety of batches. Secondly and thirdly, considerations around the definition of the baseline distribution, which provides the location and scale parameters in the preparation of response profiles, as detailed in **Materials and Methods section**

**2.4.** The determination of which type of samples to utilize for the baseline distribution was considered, whether vehicle samples (e.g., DMSO-treated) or all samples including both vehicle and treatment, and whether samples within each batch or across whole batches. Response profiles were computed under each of these 2^3^ combinations, and intraDC was subsequently evaluated.

Under the conditions with batch correction, negligible differences were evident, except for the baseline defined by vehicle control within each batch (**Figure 1**). On the other hand, in the absence of batch correction, baselines defined by samples within each batch outperformed those defined by samples across whole batches. This outcome is rational, as baselines defined by samples across whole batches are susceptible to batch differences without correction. The employment of the baseline defined by all samples within each batch, with batch correction, yielded the most favorable outcomes in the larger dataset. It is worth noting that the results for the smaller dataset (HT-HG-U133A_EA) exhibited a parallel trend while the baseline defined by all samples across whole batches with batch correction, yielding the optimal outcome. This variation can be attributed to differences in the null distribution in the calculation of intraDC across datasets (**Materials and Methods 3.1.**). It is also noteworthy that we employed MAD z for the computation of location and scale parameters, as no significant deviations were observed between z, MAD z, and robust z (**Supplementary Figure 3**).

### 2. Comparison of inter-dataset consistency of response profiles between conversion methods

**4.** Good response profiles for a compound of interest should exhibit a notable similarity to response profiles of the same compound derived from a distinct dataset. Thus, we investigated the impact of calculation methodologies on the similarity of response profiles for identical compounds originating from different sources, a metric denoted as inter-dataset consistency (interDC) in this study. InterDC is an indicator of robustness within a dataset. Holding the response profiles of the larger dataset, HT-HG-U133A, as the ground truth, established from the optimal conditions identified in the prior section, we conducted parallel investigations on the smaller dataset, HT-HG-U133A_EA, focusing on the 22 compounds shared between the two datasets. As a result, the baseline defined by all samples within each batch with batch correction, demonstrated the highest interDC, closely followed by the baseline defined by all samples across whole batches with batch correction, exhibiting a marginal difference (**Figure 2**).

**Figure 1.**
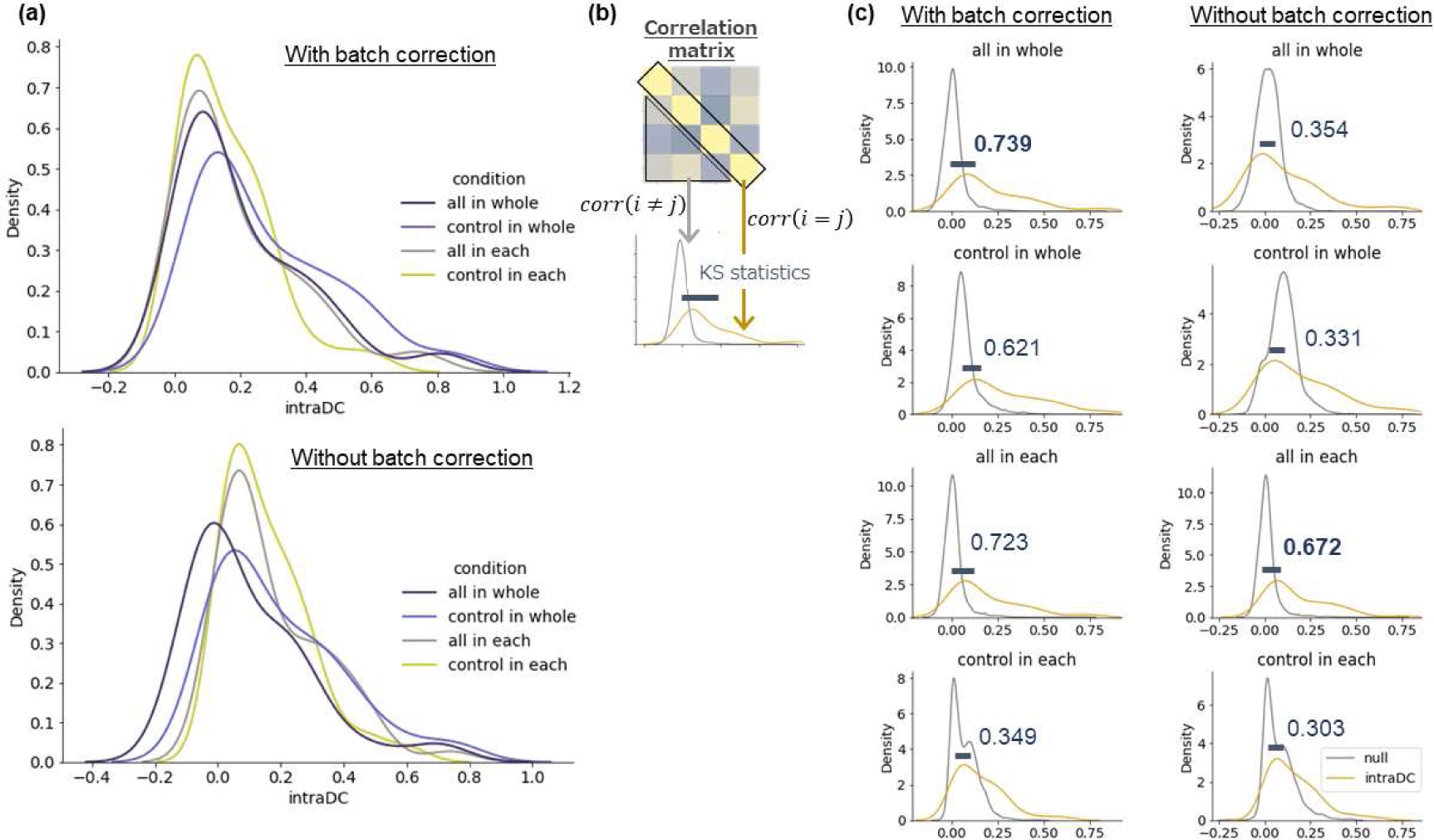
Comparison of intra-dataset consistency of response profiles between conversion methods. (a) Distribution of intra-dataset consistency (intraDC). IntraDC was calculated under the indicated conditions and histograms were plotted. The upper and lower panels show intraDC calculated from the datasets prepared with or without batch correction before conversion. all in whole, normalized by all samples in whole dataset (navy); control in whole, normalized by control samples in whole dataset (blue); all in each, normalized by all samples in each batch (grey); control in each, normalized by control samples in each batch (yellow). (b) Illustration of how to calculate intraDC and the null distribution. (c) Comparison with null distribution. Null distribution for each condition was generated and plotted with the corresponding intraDC distribution. Kolmogorov-Smirnov (KS) statistics between the intraDC and null distribution was calculated for each condition and depicted as KS with the highest value in bold.

**Figure 2.**
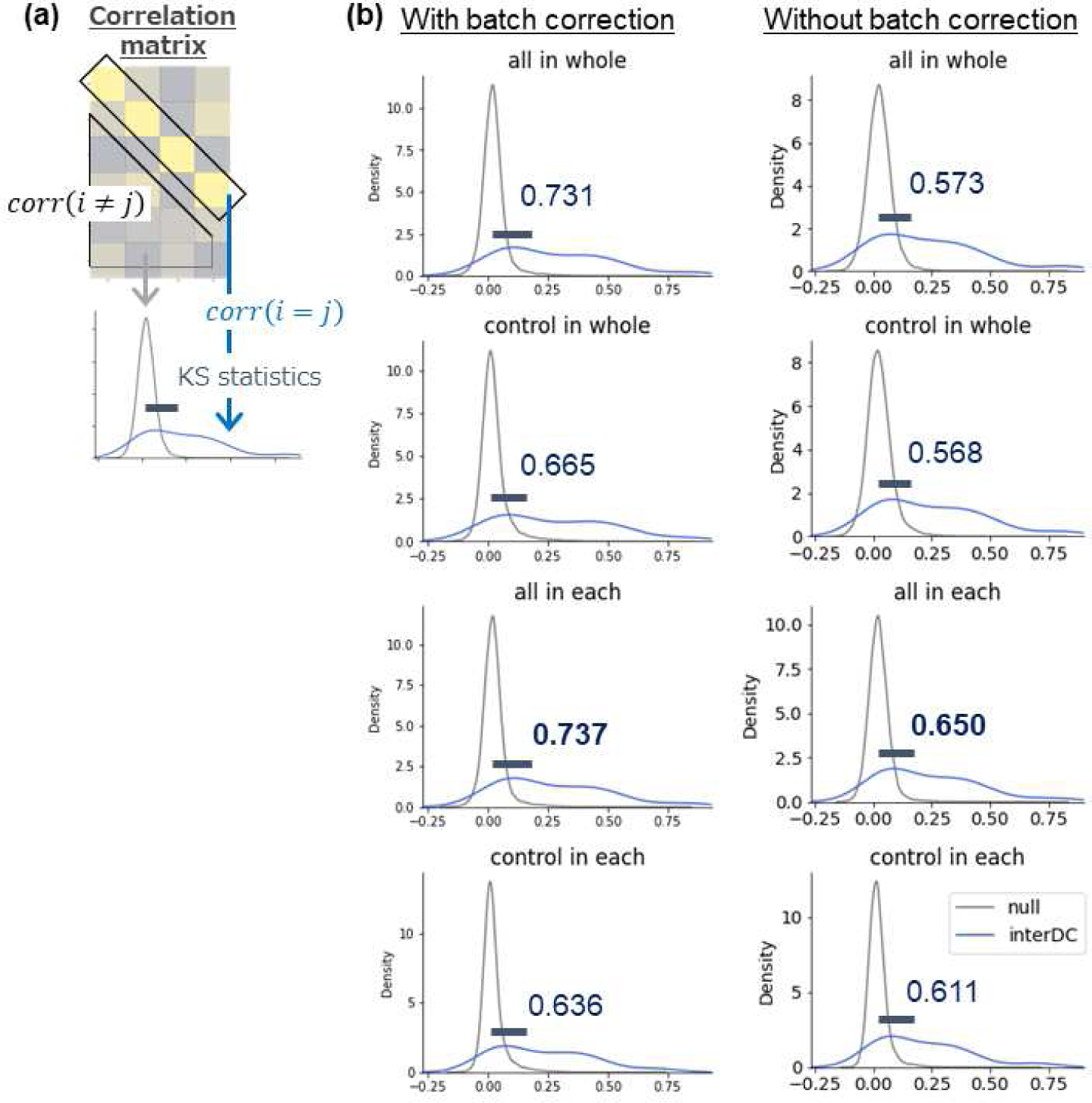
Comparison of inter-dataset consistency of response profiles between conversion methods. (a) Illustration of how to calculate inter-dataset consistency (interDC) and the null distribution. (b) Distribution of interDC compared with null distribution. Null distribution for each condition was generated and plotted with the corresponding interDC distribution. KS statistics between the interDC and null distribution was calculated for each condition and depicted as KS with the highest value in bold.

When taken into consideration on the intraDC results, these findings underscore the significance of utilizing the baselines defined by all samples with batch correction for the robust preparation of response profiles, regarding relatively large datasets. It is important to note that there existed a discrepancy between the interDC of the baseline defined by all samples within each batch and that of the baseline defined by all samples across all batches in the absence of batch correction. However, this discrepancy was largely mitigated by the application of batch correction. This suggests that batch correction effectively addresses the batch-related discrepancies that impact response profiles in the smaller dataset.

### 3. Effects of quality control process on inter-dataset consistency

As shown in **Supplementary Figure 4**, we observed a positive correlation between intraDC and interDC across various methodological variations. This implies that intraDC, which can be calculated within a dataset, serves as an indicator of interDC, which needs additional distinct dataset for calculation, rendering it a valuable tool for assessing the quality of response profiles for compounds of interest. To substantiate this point, we established *r* as the threshold for intraDC and investigated its influence on interDC. As a result, response profiles that met the *r* threshold exhibited elevated interDC as *r* increased (**Figure 3a**). Conversely, the number of compounds surpassing the threshold decreased, presenting a trade-off between the enhanced interDC and the reduced number of qualifying response profiles – a phenomenon to be expected (**Figure 3b**). These observations underscore the utility of interDC as a quality assessment metric. However, the determination of the hyperparameter *r* necessitates a careful consideration of the equilibrium between quality and availability of response profiles.

**Figure 3.**
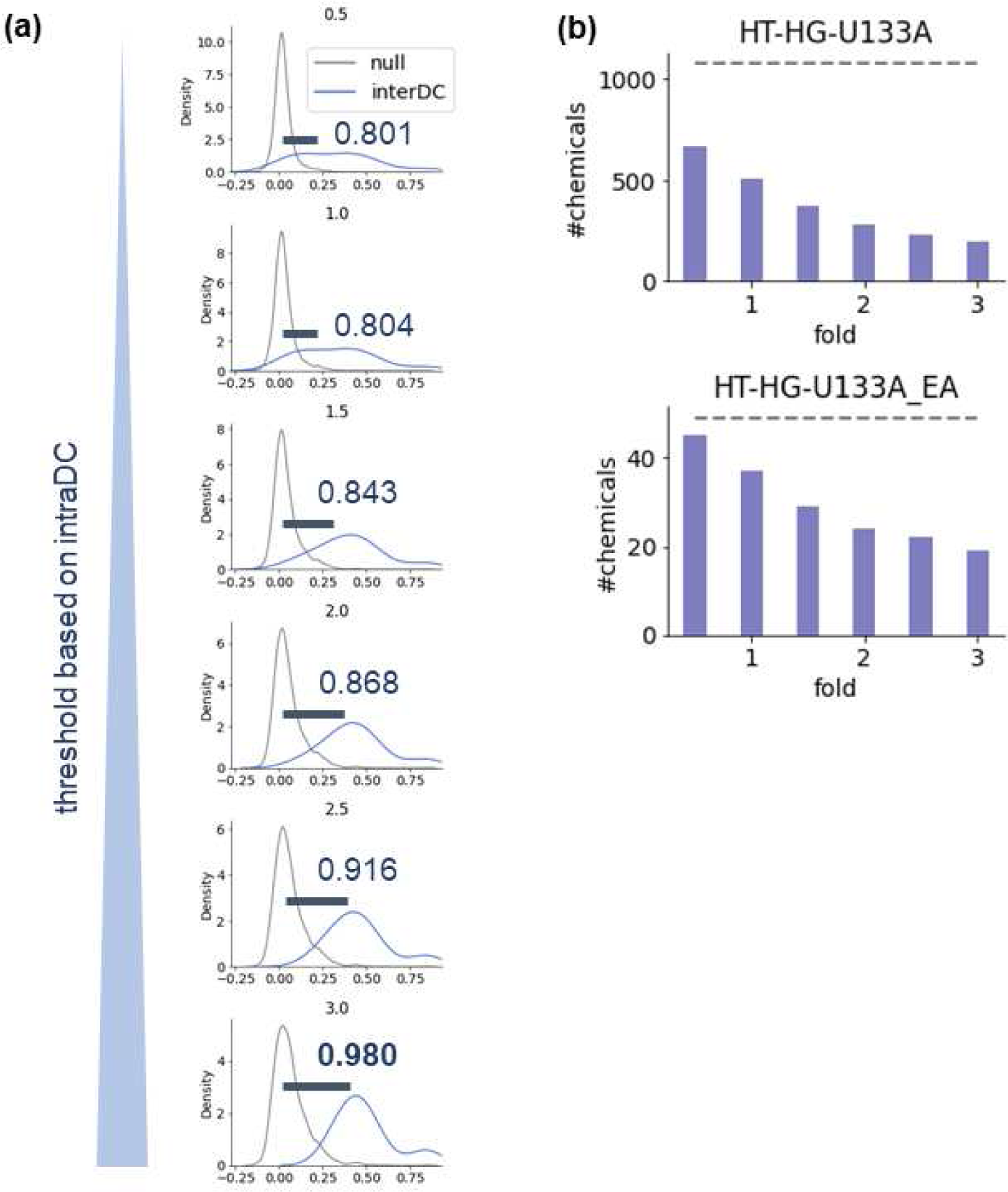
Effects of quality control process on inter-dataset consistency. (a) Relationship between the threshold of quality control and interDC. Null distribution for each condition was generated and plotted with the corresponding interDC distribution. KS statistics between the interDC and null distribution was calculated for each condition and depicted as KS with the highest value in bold. The threshold value for intraDC (*r*) is presented on the top of each panel. (b) Number of the remaining samples after quality control process with various thresholds. The grey dashed line indicates the number of compounds without quality control process.

### 4. Investigation of the number of control samples for response profiles

Until now, we have dealt with relatively large microarray datasets, with the smaller dataset HT-HG-U133A_EA alone comprising over 200 data points. However, in practical scenarios necessitating transcriptome profiles of compounds—such as in toxicity assessments and uncovering unrecognized effects—a much smaller dataset is preferable due to considerations of running costs and the number of compounds for testing. How should we design a transcriptome analysis under these situations?

From a normalization perspective, we tackled to shedding light on the above question, focusing on the baseline distribution for response profile preparation. More precisely, we conducted a simulation to investigate the influence of the number of control samples on interDC, using the response profiles from the larger dataset as the ground truth (**Figure 4a**). It is important to highlight that in this experiment, we defined the baseline distribution with vehicle samples. This choice stems from the assumption that the scope of biological responses is likely to be restricted in small datasets obtained in practical scenarios.

**Figure 4.**
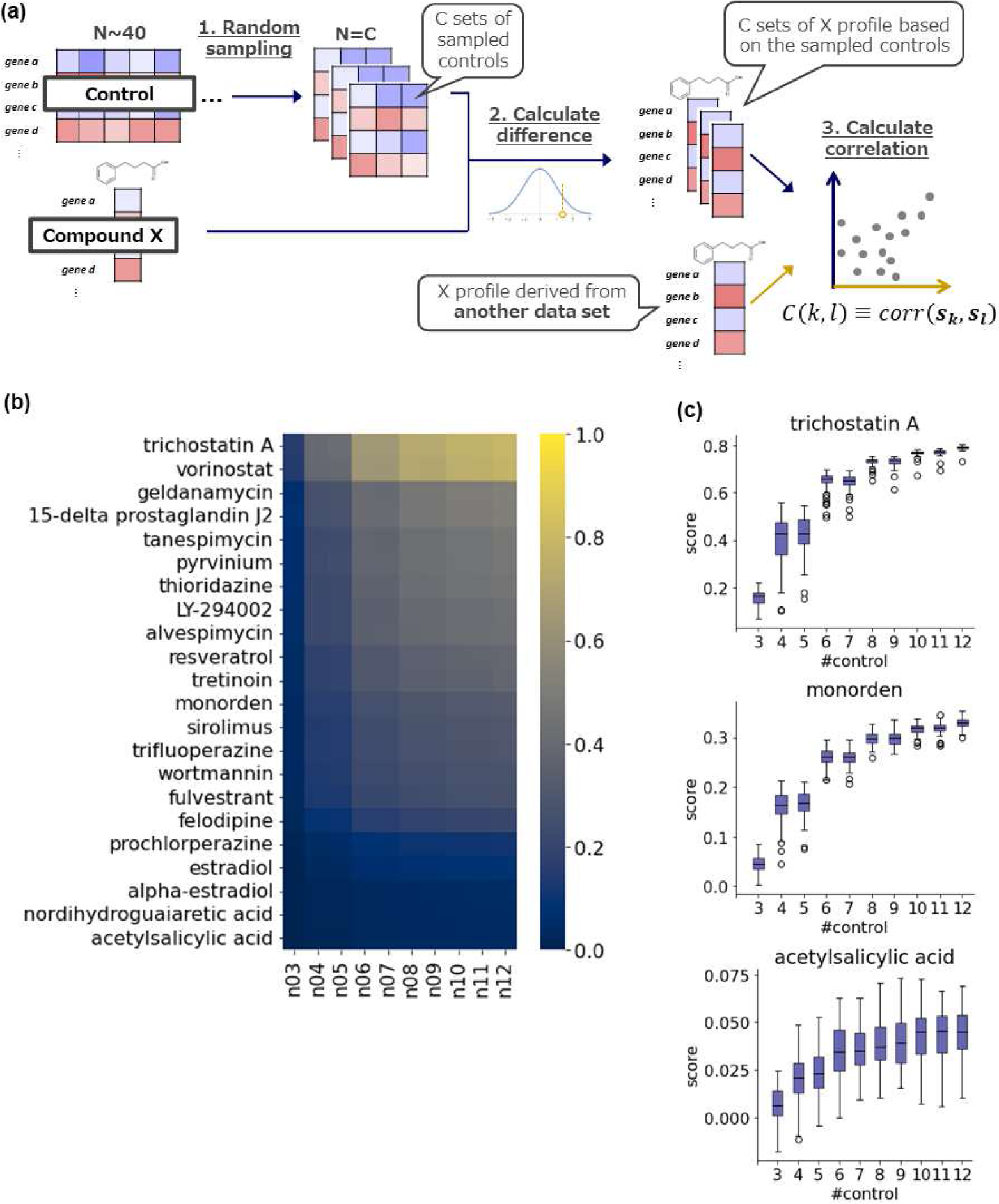
Investigation of the number of control samples for response profiles. (a) Illustration of how to evaluate the relationship between the number of control samples employed for response profile calculation and interDC. (b) Heatmap of the mean values of interDC of each compound and each number of control samples. (c) Box plots of interDC of response profiles derived from the indicated number of control samples were shown for each compound. 50 control sample sets are prepared by random sampling from the total 36 control samples in HT-HG-U133A_EA dataset and visualized. The featured compounds representing a broad spectrum of interDC are shown here and the remaining compounds are displayed in Supplementary Figure 5.

Figure 4b displays a heatmap illustrating the mean values of interDC for the 22 compounds in relation to the number of control samples. A global trend emerged, wherein interDC exhibited a concurrent rise along with the increase in the number of control samples. Upon visualizing the relationship between interDC and the number of control samples for each compound, it became apparent that all compounds demonstrated a consistent upward trajectory in interDC, irrespective of their initial interDC scores (Figure 4c **and Supplementary Figure 5**). This pattern suggests that the number of control samples should ideally be 6 or greater. Theoretically, this would be comprehensible as an unbiased variance estimate derived from a Gaussian distribution adheres to a chi-square distribution if inter-DC conforms to a Gaussian distribution. **Supplementary Figure 6** illustrates the 95% confidence intervals of standard deviation estimates of standard normal distribution based on the specified number of samples. The confidence intervals tend to stabilize around 5 or 6, mirroring the simulation results in Figure 4c.

## Conclusion

The contributions of this study are as follows:

1. In calculation of response profiles, the baseline distribution defined by all samples within each batch, with batch correction, is a good choice for handling large datasets (n > 100, in our opinion).
2. We have demonstrated a positive correlation between consistency intra-dataset and inter-dataset, indicating that intra-dataset consistency can serve as a quality check for response profiles.
3. We propose that the number of control samples should ideally be 6 or greater in small datasets.

A notable limitation of this study is that all the results are derived from only two microarray datasets *in vitro*. Given the prevailing trend, the exploration utilizing RNA-seq data must be undertaken within the realm of practical applicability, even though benchmark datasets of suitable scale are presently unavailable. It is also noteworthy that determining the appropriate sample size for acquiring *in vivo* transcriptome profiles is crucial as well, aligning with the principles of the 3R. While it is challenging to assemble additional datasets of a similar scale, further validation is imperative to ascertain the generalizability of these findings. It is essential to acknowledge that certain compounds exhibited markedly low inter-dataset consistency. Compounds demonstrating high interDC, with robust response profiles across datasets, appear to exert potent perturbations on cells, such as HDAC inhibitors and anti-cancer agents, in contrast to compounds with low inter-dataset consistency, such as acetylsalicylic acid. The presence of a systematic trend classifying compounds into high or low inter-dataset consistency categories is an intriguing question, which could be also addressed through the expansion of evaluation datasets and subsequent validation. We believe that this study contributes to a better understanding of how to leverage transcriptome profiles of compounds and encourages the accumulation of knowledge essential for the practical utilization of such profiles, including their applicability.

## Acknowledgement

We thank all those who contributed to the construction of the datasets employed in the present study such as CMap. This work was supported by a grant-in-aid of Mochida memorial foundation for medical and pharmaceutical research.

## Author’s contributions

Tadahaya Mizuno: Conceptualization, Resources, Supervision, Project administration, Funding acquisition, Methodology, Software, Investigation, Visualization, Writing – Original Draft, Writing – Review & Editing.

Hiroyuki Kusuhara: Writing – Review & Editing

## Supplementary Figures and Tables

**Supplementary Figure 1.**
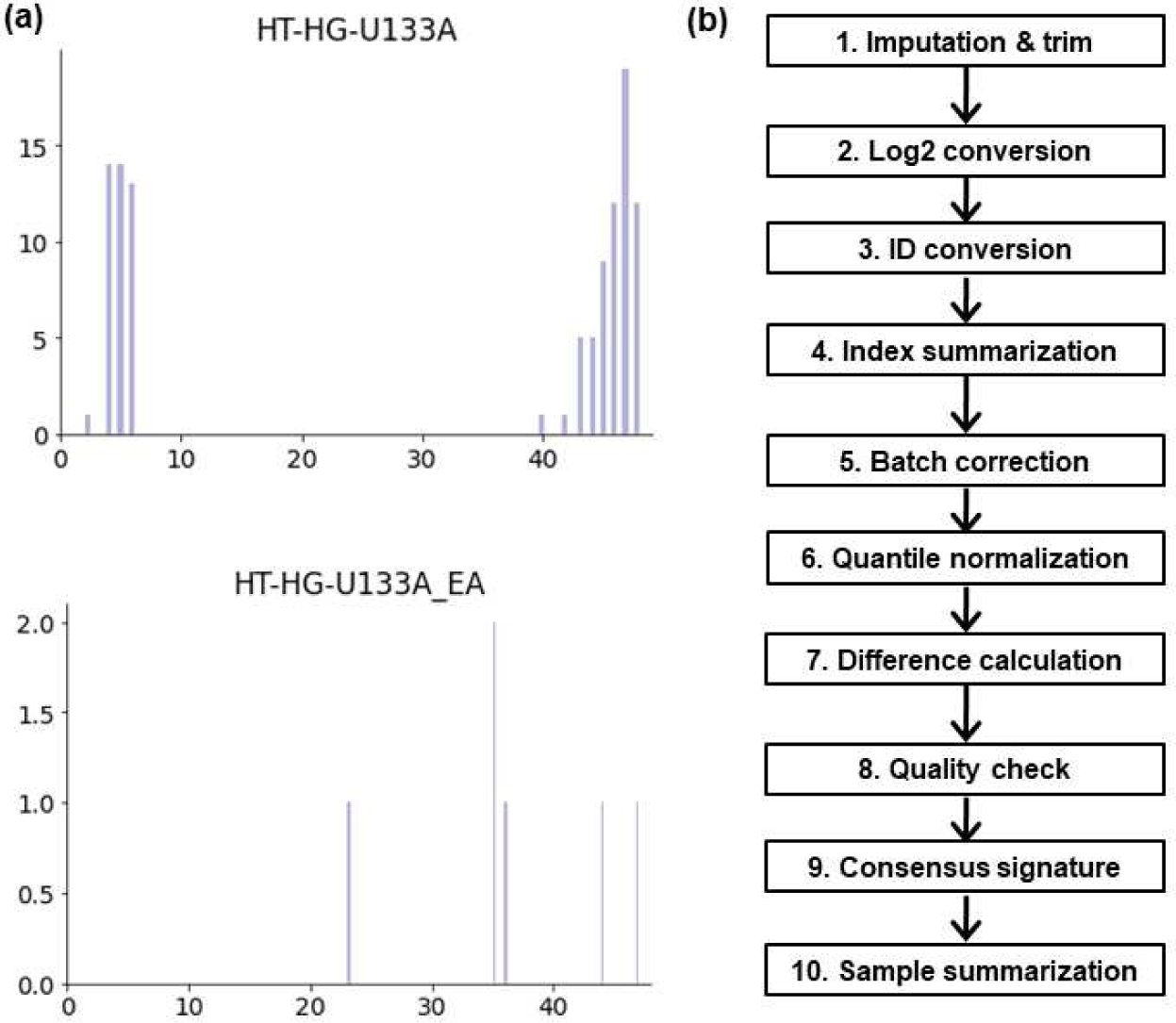
Data information and processing flow employed in this study. (a) Histogram of the number of samples in each batch of CMap datasets (HT-HG-U133A and HT-HG-U133A_EA). (b) Flow chart of expression to response. This processing flow is summarized in *exp2res* package in python 3.

**Supplementary Figure 2.**
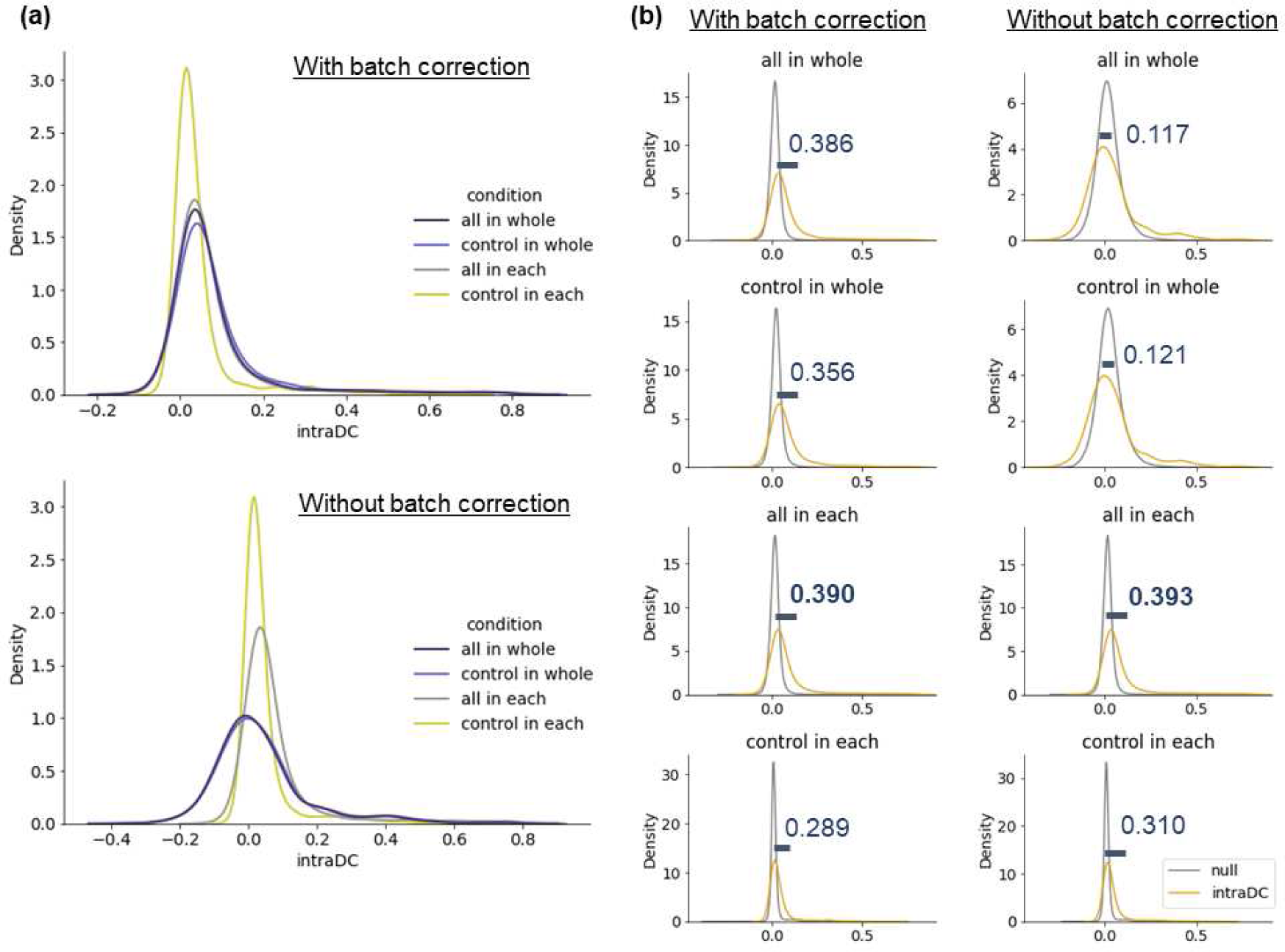
Comparison of intra-dataset consistency of response profiles between conversion methods with HT-HG-U133A dataset. (a) Distribution of intra-dataset consistency (intraDC). IntraDC was calculated under the indicated conditions and histograms were plotted. The upper and lower panels show intraDC calculated from the datasets prepared with or without batch correction before conversion. all in whole, normalized by all samples in whole dataset (navy); control in whole, normalized by control samples in whole dataset (blue); all in each, normalized by all samples in each batch (grey); control in each, normalized by control samples in each batch (yellow). (b) Comparison with null distribution. Null distribution for each condition was generated and plotted with the corresponding intraDC distribution. Kolmogorov-Smirnov statistics between the intraDC and null distribution was calculated for each condition and depicted as KS with the highest value in bold.

**Supplementary Figure 3.**
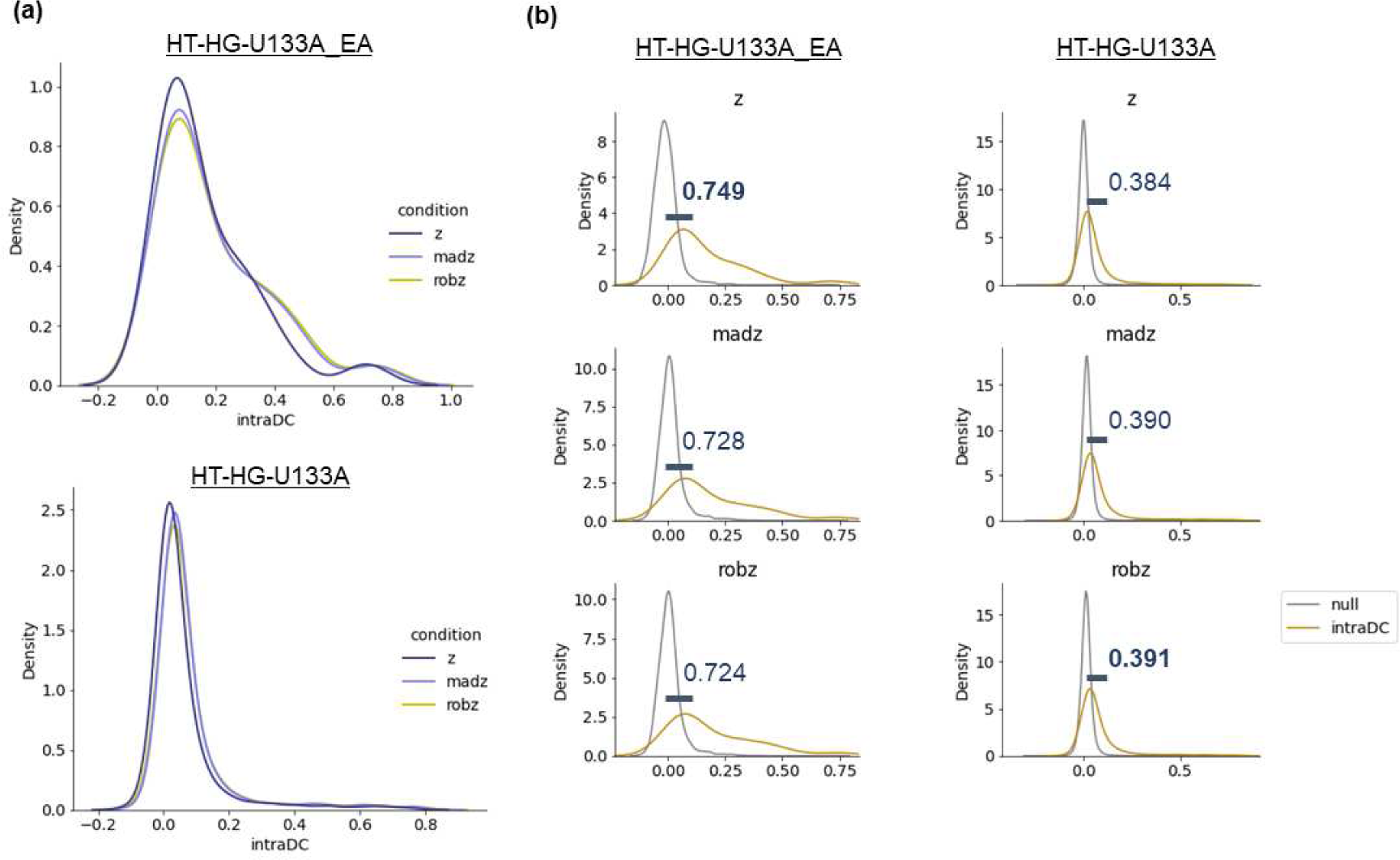
Comparison between z, modified z, and robust z scores in response calculation. (a) Distribution of intra-dataset consistency (intraDC). IntraDC was calculated under the indicated conditions and histograms were plotted. The upper and lower panels show intraDC calculated from HT-HG-U133A_EA and HT-HG-U133A datasets. z, z score (navy); madz, modified z score (blue); robz, robust z score (yellow). (b) Comparison with null distribution. Null distribution for each condition was generated and plotted with the corresponding intraDC distribution. Kolmogorov-Smirnov statistics between the intraDC and null distribution was calculated for each condition and depicted as KS with the highest value in bold.

**Supplementary Figure 4.**
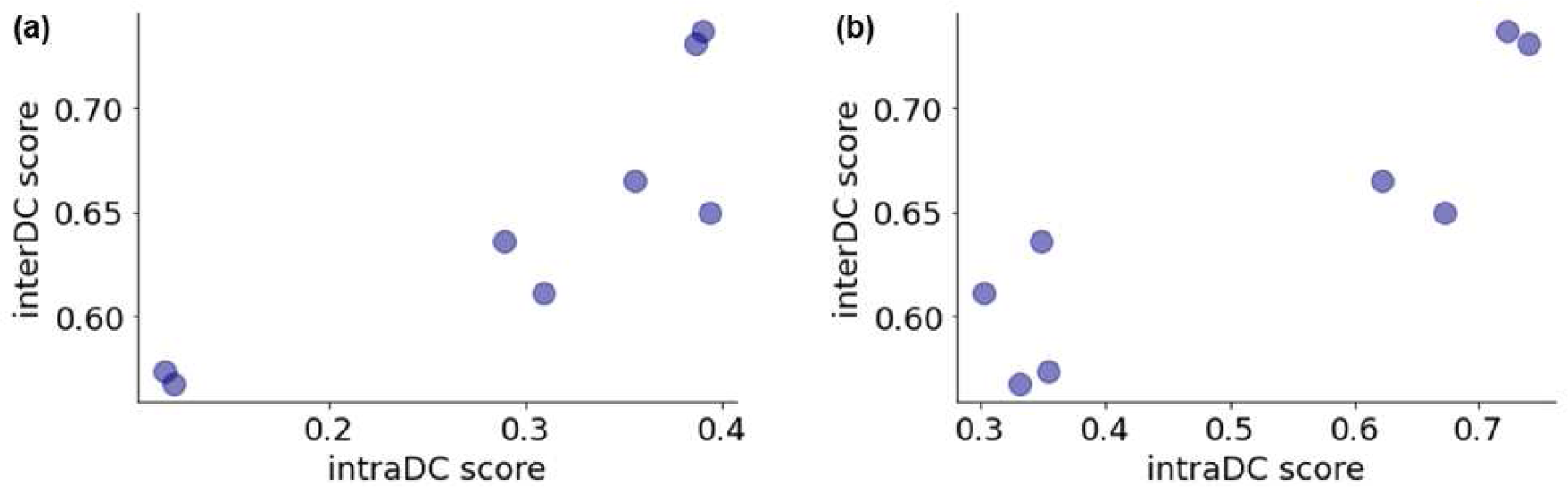
Correlation between intraDC and interDC scores. (a) and (b) show scatter plot between interDC and intraDC scores of HT-HG-U133A and HT-HG-U133A_EA datasets obtained in Results and Discussion section 1 and 2, respectively.

**Supplementary Figure 5.**
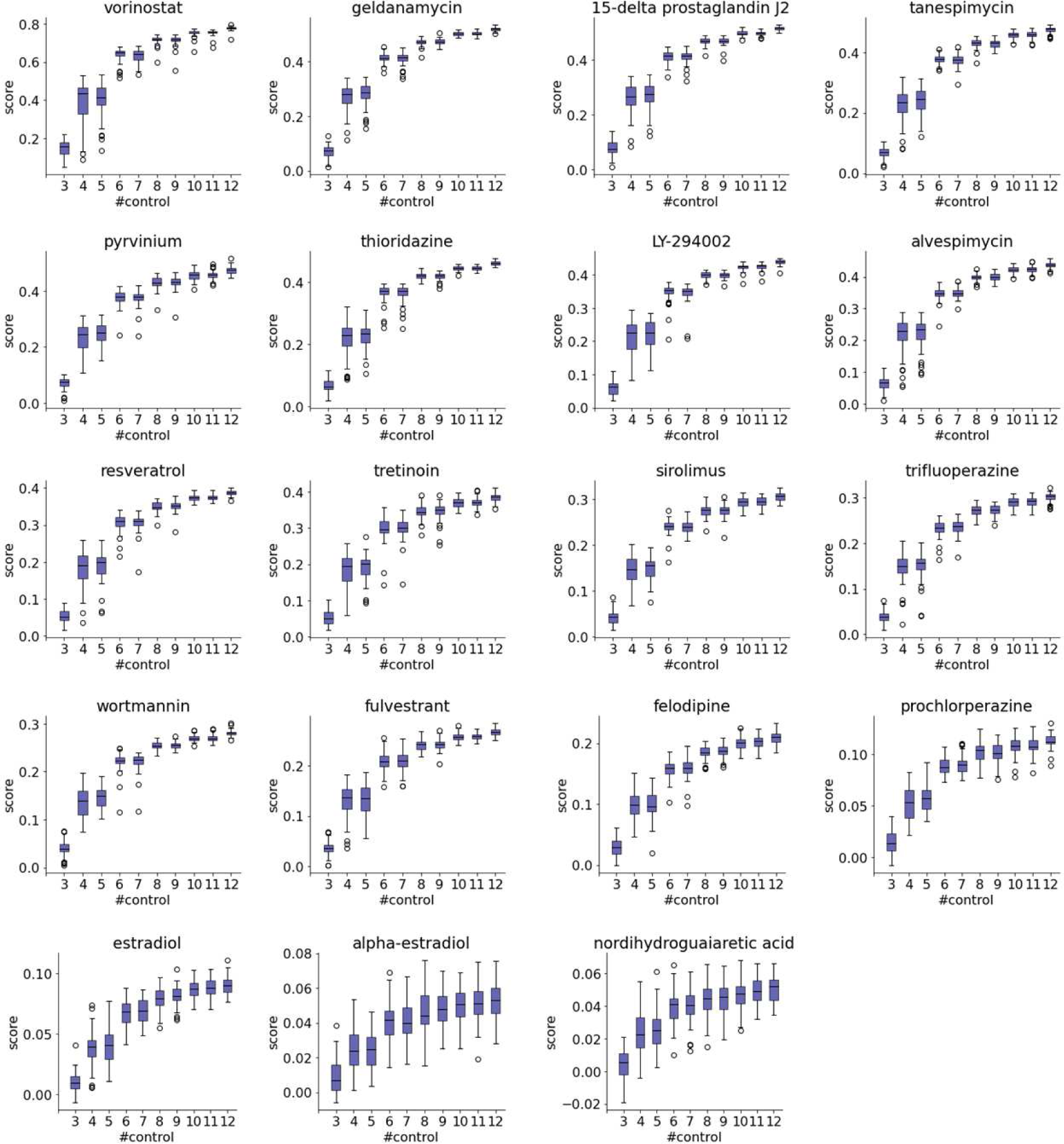
Relationship between the number of control samples and interDC. Box plots of interDC of response profiles derived from the indicated number of control samples were shown for each compound. 50 control sample sets are prepared by random sampling from the total 36 control samples in HT-HG-U133A_EA dataset and visualized. The compounds not shown in the main text are paneled here.

**Supplementary Figure 6.**
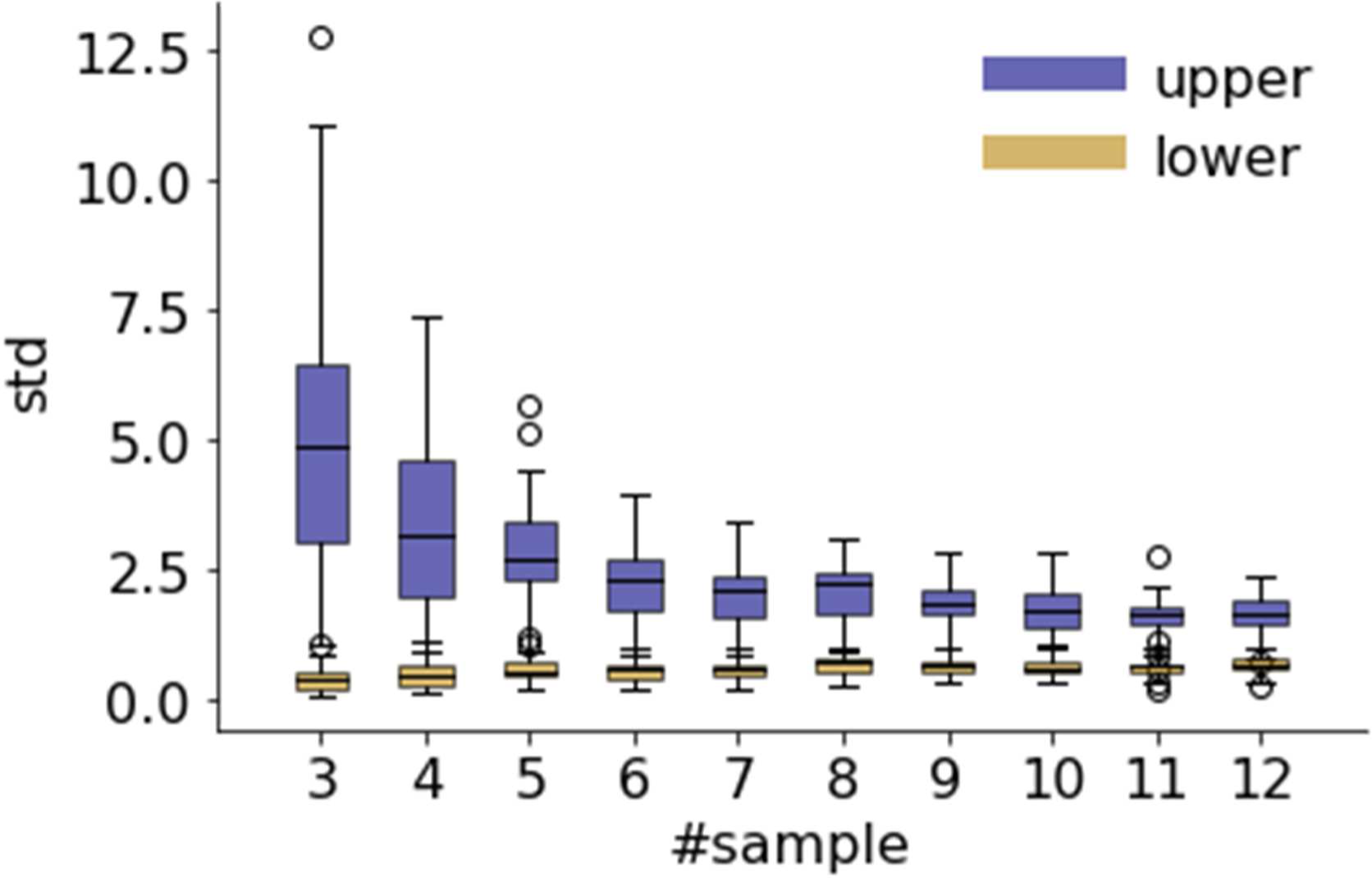
Relationship between the number of samples and stability of standard deviation. Box plots of the upper and lower 95% confidential intervals (CI) of standard deviation estimated by the indicated number of random normal. In the scenario where the population follows to a normal distribution with a population variance of *a-^2^*, and samples of size *n* are drawn from it resulting in an unbiased variance of *s^2^*, the ensuing *x^2^* statistic adheres to a chi-square distribution with *n − 1* degrees of freedom. The indicated number of samples are drawn from standard normal distribution for 50 times and standard deviation is estimated. X-axis and Y-axis indicate the number of random normal utilized for estimation and standard deviation, respectively. The upper and lower CI are depicted as navy and gold, respectively.

**Supplementary Table 1.** Concentration of compounds employed for investigating inter-dataset consistency.

| **Name** | **Concentration (μM)** |
| --- | --- |
| 15-delta prostaglandin J2 | 10 |
| acetylsalicylic acid | 100 |
| alpha-estradiol | 0.01 |
| alvespimycin | 0.1 |
| estradiol | 0.1 |
| felodipine | 10 |
| fulvestrant | 1 |
| geldanamycin | 1 |
| LY-294002 | 10 |
| monorden | 0.1 |
| nordihydroguaiaretic acid | 1 |
| prochlorperazine | 10 |
| pyrvinium | 1.25 |
| resveratrol | 10 |
| sirolimus | 0.1 |
| tanespimycin | 1 |
| thioridazine | 10 |
| tretinoin | 1 |
| trichostatin A | 1 |
| trifluoperazine | 10 |
| vorinostat | 10 |
| wortmannin | 1 |

